# The Sin3B chromatin modifier restricts cell cycle progression to dictate hematopoietic stem cell differentiation

**DOI:** 10.1101/2023.01.23.525185

**Authors:** Alexander Calderon, Tamara Mestvirishvili, Francesco Boccalatte, Kelly V. Ruggles, Gregory David

## Abstract

To maintain blood homeostasis, millions of terminally differentiated effector cells are produced every day. At the apex of this massive and constant blood production lie hematopoietic stem cells (HSCs), a rare cell type harboring unique self-renewal and multipotent properties. A key feature of HSCs is their ability to temporarily exit the cell cycle in a state termed quiescence. Defective control of cell cycle progression can eventually lead to bone marrow failure or malignant transformation. It is thought that HSCs must re-enter the cell cycle in order to commit to terminal differentiation. However, the molecular mechanisms tying cell cycle re-entry to cell fate commitment in HSCs remain elusive. Here, we identify the chromatin-associated Sin3B protein as a molecular link between cell cycle progression and differentiation in HSCs. We demonstrate that Sin3B is necessary for HSCs’ commitment to differentiation, but dispensable for their self-renewal or survival. Single cell transcriptional profiling of hematopoietic stem and progenitor cells (HSPCs) inactivated for Sin3B reveals aberrant cell cycle gene expression, consistent with the observed aberrant progression through the G_1_ phase of the cell cycle. The defective cell cycle control elicited upon Sin3B inactivation correlates with the engagement of discrete signaling programs, including aberrant expression of cell adhesion molecules and essential components of the interferon signaling cascade in LT-HSCs. Additionally, chromatin accessibility profiling in LT-HSCs reveals the Sin3B-dependent accessibility of genomic elements controlling HSC differentiation, suggesting a functional link between cell cycle progression, and priming of hematopoietic stem cells for differentiation. Together, these results point to controlled progression through the G_1_ phase of the cell cycle as a likely regulator of HSC lineage commitment through the modulation of chromatin features.

## Introduction

Hematopoietic stem cells (HSCs) are rare, multipotent, self-renewing adult stem cells responsible for maintaining hematopoiesis throughout the lifespan of an organism^1^. HSCs sit at the top of a hierarchically organized system of progenitors that give rise to terminally differentiated effector blood cells with distinct roles^2^. To maintain their stemness features, both extrinsic and intrinsic factors enforce a quiescent state in HSCs^3^. Quiescence is a form of temporary cell cycle exit and is thought to support the long-term fitness of HSCs by shielding them from environmental damage and cellular injury generated through metabolic processes^4^. Extrinsic factors regulating HSC quiescence stem from their unique localization within specialized regions of the bone marrow called niches. Intrinsic factors dictating HSC quiescence include canonical cell cycle pathways, such as Rb-E2F repression.

The proliferative status of HSCs inversely correlates with their functional potency, as actively cycling cells exhibit diminished reconstitution potential when transplanted into lethally irradiated mice^5^. Additionally, label retention studies revealed that a subpopulation of dormant, *i*.*e*. non-proliferative, HSCs encompasses the entire reconstitution capacity of the stem cell compartment^6^. When hematopoietic demand increases, these dormant HSCs can enter a primed or activated state that potentiates faster re-entry into the cell cycle to re-establish homeostasis^7,8^. Activated HSCs may also revert to a dormant state, but the molecular mechanisms regulating this reversible switch have not been fully elucidated.

The contribution of cell cycle phases in regulating differentiation of some non-hematopoietic stem cells, namely embryonic stem cells (ES), has been well documented^9^. ES cells possess a unique cell cycle structure with a truncated G_1_ phase^10^. This short G_1_ phase is thought to promote self-renewal of ES cells by limiting their exposure to pro-differentiation signals when they are most susceptible to the engagement of differentiation programs^11^. Accordingly, lengthening the G_1_ phase causes spurious differentiation in ES cells^12^. Additionally, as ES cells normally differentiate, their G_1_ transit time through G_1_ also increases^13^. However, whether location in a discrete cell cycle phase is a cause or consequence of differentiation in ES cells remains unresolved.

The relationship between cell cycle and differentiation in adult stem cells remains even more ill-defined. HSCs, like other somatic cells, possess a substantial G_1_ phase subject to regulation by the canonical cell cycle machinery. A regulatory function for G_1_ in HSC differentiation has been proposed, notably based on the observation that CCND1-CDK4 overexpression in human HSCs shortened G_1_ and biased cellular fate towards self-renewal at the expense of differentiation^14^. This hypothesis highlighting a role for G_1_ is further supported by a study demonstrating that HSCs can differentiate without undergoing mitosis, suggesting that cellular division is dispensable for differentiation in HSCs^15^. Finally, a recent report has shown that the relative speed of cell cycle progression in megakaryocytic-erythroid progenitors (MEPs) determines their fate specification, as faster cell cycle progression promotes the erythroid fate over the megakaryocytic fate^16^. Together, these data suggest that cell cycle progression *per se* could serve as a determinant of differentiation in the hematopoietic system. However, how cell cycle machinery affects transcriptional output related to differentiation processes in HSCs remains unknown.

To begin to dissect the relationship between cell cycle progression in G_1_ and hematopoietic stem cell differentiation, we exploited a mouse strain where the transcriptional repressor Sin3B can be somatically inactivated in the hematopoietic system^17^. Sin3B is an evolutionarily conserved non-catalytic scaffolding protein that is part of the Sin3 transcriptional repressor complex^18^. Through its interaction with sequence-specific transcription factors, Sin3B tethers histone deacetylase (HDAC) activity to discrete genomic loci. We have shown that Sin3B is a key contributor to cell cycle exit in numerous biological contexts^19–21^. Additionally, we have demonstrated that Sin3B partially mediates the repressive activity of the <u>D</u>imerization Partner (DP), <u>R</u>etinoblastoma (RB)-like, <u>E</u>2F, <u>a</u>nd <u>M</u>uvB (DREAM) complex, another critical regulator of quiescence and G_1_ entry^22^. Genetic inactivation of Sin3B in the hematopoietic system via the Vav1-iCre transgene impairs HSCs quiescence and abolishes their ability to reconstitute the hematopoietic system in a competitive transplantation setting but does not affect their self-renewal and survival^17^. Thus, Sin3B inactivation uncouples self-renewal and differentiation in HSCs and offers a unique opportunity to study the relationship between cell cycle progression and differentiation in the adult stem cell compartment.

Accordingly, loss of Sin3B results in spurious progression through the early phases of the cell cycle in HSCs, but single cell transcriptome analysis reveals only nominal changes in the expression of non cell cycle-related genes. Whether cell cycle location and differentiation are linked similarly in HSCs and their progeny remain unknown. Our results identify Sin3B as both an essential gatekeeper of early cell cycle progression in HSCs, and a molecular switch for HSC lineage commitment through the modulation of chromatin accessibility at cis-regulatory elements driving hematopoietic differentiation. Together, these results point to controlled progression through the G_1_ phase of the cell cycle as a potential regulator of HSC lineage commitment through the modulation of chromatin features.

## Results

### Genetic inactivation of Sin3B results in LT-HSC expansion and aberrant transcriptional signatures

We have previously demonstrated that Sin3B^-/-^ whole bone marrow cells (WBM) are unable to reconstitute the hematopoietic systems of lethally irradiated mice in a competitive transplantation setting^17^. Peripheral blood analysis of recipient mice over 20 weeks showed minimal contribution of Sin3B^-/-^ cells to all hematopoietic lineages assayed via flow cytometry. At 20 weeks, when the hematopoietic system has returned to homeostasis, we analyzed the bone marrow to determine the contribution of donor cells to the stem cell compartment. Surprisingly, the analysis revealed that Sin3B^-/-^ Long Term (LT)-HSCs are present in the bone marrow in comparable proportions to their wild-type counterparts, despite minimal contributions to the peripheral blood. However, these experiments did not reveal the contribution of Sin3B^-/-^ hematopoietic cells to the stem cell compartment throughout a stress response. In particular, whether Sin3B^-/-^ LT-HSCs are blocked in their differentiation at the niche, or if their differentiation is merely delayed until later timepoints is unknown. To better characterize the differentiation defects elicited upon Sin3B inactivation, we analyzed the bone marrow compartment of lethally irradiated recipient mice 8 weeks post transplantation, when LT-HSCs are actively engaged in the stress response. Using a panel of markers including SLAM receptors^23^, we documented the proportion of different hematopoietic stem and progenitor cells in the bone marrow of these recipient mice. Sin3B^-/-^ WBM contributed minimally to total bone marrow cellularity (16%) compared to their wild-type counterparts (67%) (Figure 1A). This pattern held in more primitive hematopoietic subtypes (LSK, MPP2, MPP3, MPP4, ST-HSCs). Given the limited self-renewal capacity of MPPs in a transplantation setting, with the majority of new MPPs derived from LT-HSCs at later timepoints, we expect all mature hematopoietic stem cells at this point to originate from LT-HSCs^24^. By contrast, both wild-type and Sin3B^-/-^ bone marrow cells contributed to the LT-HSC compartment to a comparable extent (Figure 1A). This suggests that Sin3B^-/-^ LT-HSCs are present in their niche but are blocked in differentiation, even to other primitive cell types such as ST-HSCs and MPPs. Together, these data indicate that Sin3B^-/-^ LT-HSCs are able to survive, proliferate, and self-renew in a transplantation setting, but are severely impaired in their ability to differentiate.

**Figure 1.**
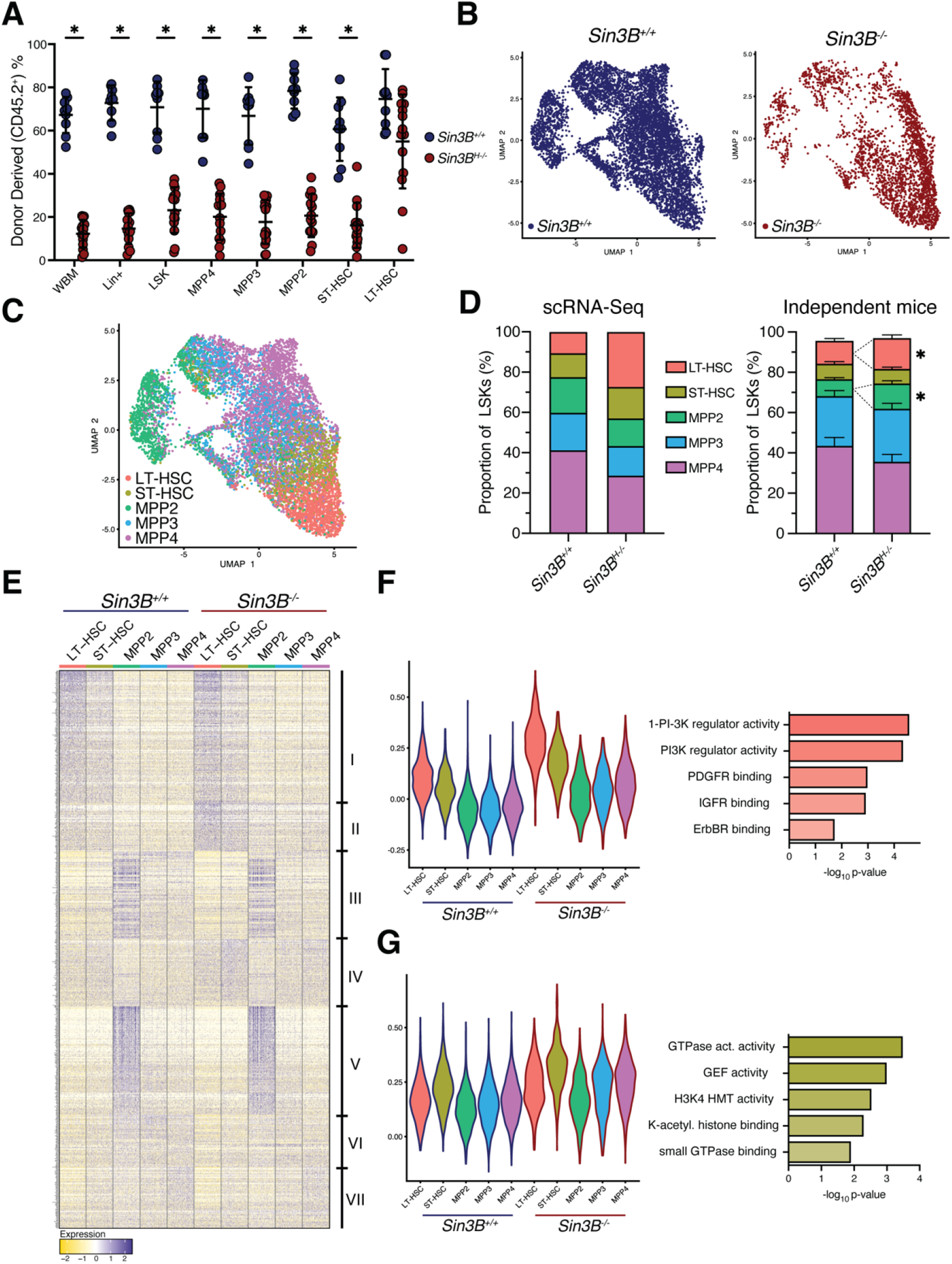
Sin3B regulates discrete transcriptional programs in hematopoietic subsets at homeostasis. **A**. Analysis of whole bone marrow from recipient mice 8 weeks after competitive transplantation. Quantification of donor derived (CD45.2) cells in the hematopoietic compartments indicated via flow cytometry. Asterisks indicate statistical significance. (2way ANOVA, S □ idák’s multiple comparison’s test; *p<0.0001; Data are represented as mean ± SEM) Sin3B^F/F^ recipients n = 9; Sin3B^-/-^ recipients n = 14. **B**. Two-dimensional Uniform Manifold Approximation and Projection (2D UMAP) with LSK cells colored by genotype as determined by hashtag oligo analysis. Left: Sin3B^+/+^ cells. Right: Sin3B^-/-^ cells. **C**. 2D UMAP) demonstrating supervised clustering of hematopoietic subsets recovered from scRNA-Seq. **D**. Left panel: quantification of various LSK subsets. Right panel: Quantification of same subset in independent mice. Asterisks indicate statistical significance (Student’s t-test with multiple comparison correction; *p<0.05; data shown as mean ± SEM) (n=9). **E**. Differentially expressed genes between all cell types was calculated via Seurat’s FindAllMarkers function utilizing the MAST statistical framework. Clusters of genes visually identified are denoted on the right. Expression is scaled and centered. **F**. Left panel: Violin Plot showing average expression of Cluster II genes for indicated hematopoietic subtypes. Right panel: Gene Ontology Analysis of Cluster II genes. **G**. Left panel: Violin Plot showing average expression of Cluster IV genes for indicated hematopoietic subtypes. Right panel: Gene Ontology Analysis of Cluster IV genes.

In order to understand the molecular basis underpinning the differentiation defect in Sin3B^-/-^ LT-HSCs, we performed single-cell RNA-Seq on the Lineage^-^Sca-1^+^cKit^+^ (LSK) compartment from Sin3B^+/+^ and Sin3B^H-/-^ mice (*Sin3B*^*F/F*^; *Vav1-iCre*^*+*^). After demultiplexing the data and performing quality control, we obtained data for 9586 cells. Utilizing hashtag oligonucleotides, we concomitantly sequenced Sin3B^+/+^ and Sin3B^-/-^ LSKs and were able to assign a genotype to each cell (Figure 1B)^25^. We subjected our dataset to supervised clustering and assigned each cell to a discrete hematopoietic subset using previously published transcriptional signatures^24,26^ (Figure 1C). We recovered all five hematopoietic subsets (LT-HSC, ST-HSC, MPP2, MPP3, MPP4) in both genotypes, consistent with the lack of an overt hematopoietic phenotype in Sin3B^H-/-^ mice at homeostasis^17^. We however noted an increase in the proportion of LT-HSCs in the LSK compartment of Sin3B^H-/-^ mice (27% vs 10%) (Figure 1D, left panel), and we experimentally confirmed this result in independent mice (Figure 1D, right panel). These results indicate that the single cell transcriptome analysis recapitulates the expansion of the LT-HSC compartment in Sin3B^H-/-^ mice at homeostasis.

Next, we sought to identify the transcriptional programs affected by the loss of Sin3B in each hematopoietic subtype. We identified 842 differentially expressed genes that we broadly grouped into 7 clusters based on expression patterns across genotypes, cell identity, or both (Figure 1E). We highlight two clusters of interest: Cluster II contains 74 genes that display the highest expression in LT-HSCs and are uniformly upregulated as a consequence of Sin3B loss in LSKs (Figure 1F, left panel). Gene Ontology analysis of these genes points to PI3K regulation. Importantly, previous reports had highlighted the role PI3K signaling as a mediator of HSC activation upon stress^27^. Cluster IV contains 102 genes that are enriched in GTPase activity (Figure 1G), with Rho GTPases long having been described as important in HSC function^28^. These genes are upregulated in Sin3B^-/-^ cells across different cell identity, with the highest expression in ST-HSCs, possibly highlighting a role in the LT- to ST-HSC transition. In particular, the Rho GTPase Cdc42 has recently been implicated in regulating symmetric vs asymmetric divisions in LT-HSCs^29^.

### A Sin3B-dependent transcriptional program is engaged upon the transition from LT- to ST-HSCs at homeostasis

We next leveraged the scRNA-Seq data we generated by analyzing how the differentiation program from LT-HSC to MPPs is altered upon Sin3B loss. To this end, we subjected the dataset to pseudotime analysis using the Monocle3 package^30^. We separated the cells by genotype to capture the differentiation trajectory in Sin3B^+/+^ cells utilizing an unsupervised algorithm that orders cells along a path, grouping cells locally by similarity in their transcriptomic profiles (Figure 2A, left panel). This procedure captures transcriptional paths that cells take as they transition between one cell type to another. The interconnectedness of the cells is reminiscent of a report that analyzed human HSPCs and put forward the idea of CLOUD-HSPCs^31^, in that stem and progenitor cells exist along a continuum of low-priming and gradually gain lineage commitment. The same analysis performed on Sin3B^-/-^ LSKs (Figure 2A, right panel) revealed a decrease in the number of trajectories, possibly denoting fewer possibilities for differentiation in the absence of Sin3B.

**Figure 2.**
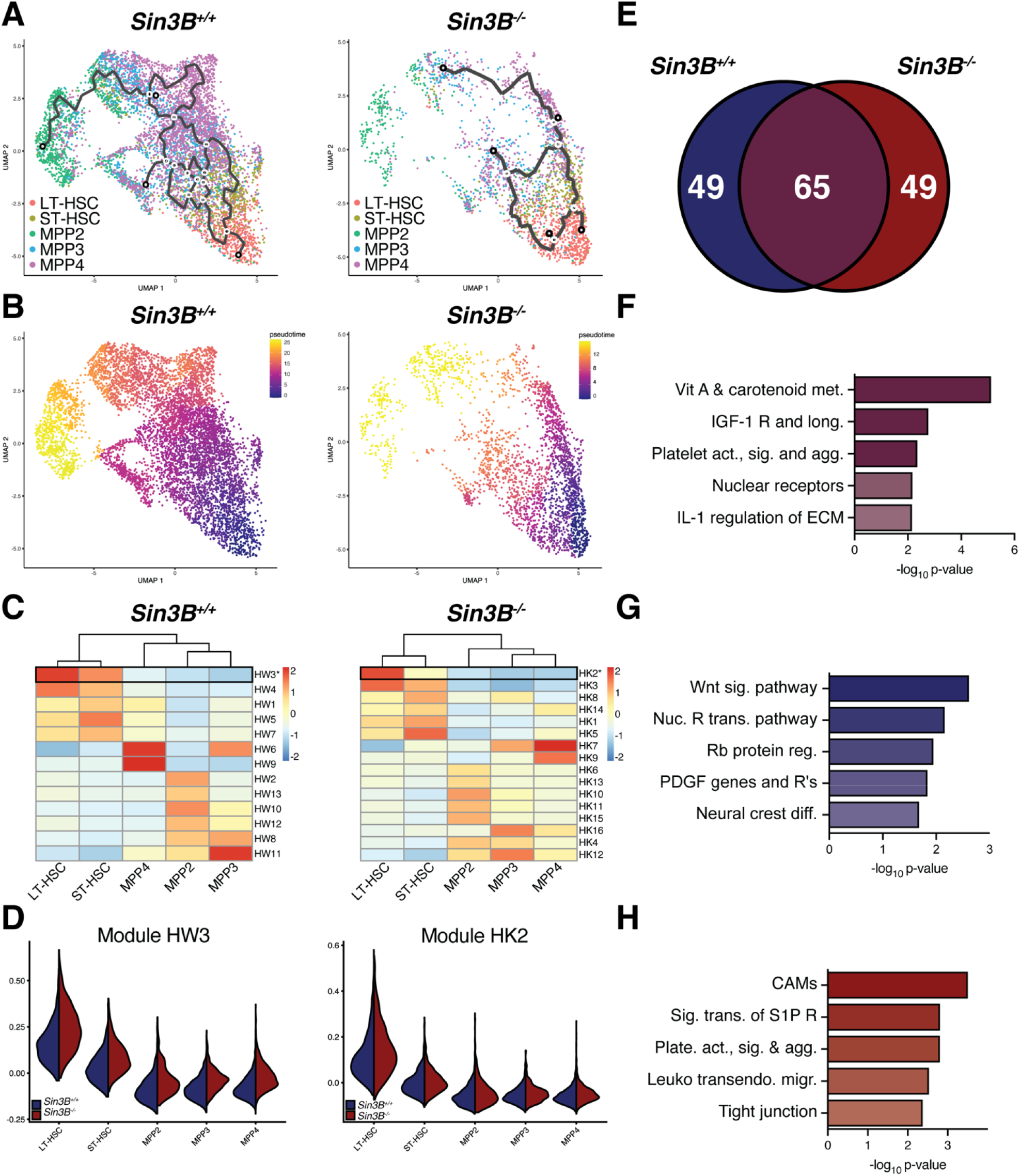
Pseudotime analysis of LSKs at homeostasis demonstrates a defective transition between LT- and ST-HSCs in the absence of Sin3B. **A**. Two-dimensional UMAP projection of LSKs showing trajectories calculated from Monocle3. Left panel: Sin3B^+/+^, Right panel: Sin3B^-/-^. **B**. Using LT-HSCs as a starting node, pseudotime was calculated for each genotype to determine transcriptional programs as a function of differentiation. **C**. Differentially expressed genes as a function of pseudotime were calculated and grouped into modules based on expression pattern. Shown is the average expression of each module in the indicated cell type. **D**. Expression of highlighted modules in Sin3B^+/+^ and Sin3B^-/-^ subsets. **E**. Venn diagram showing overlap of genes obtained from module HW3 and HK2. Purple color denotes overlap, blue denotes wild-type, and red denotes knockout. **F**. Gene Ontology Molecular Function was carried out for the indicated gene list and shown are GO terms recovered plotted by p-value. **G**. GO Molecular Function was carried out for 49 genes unique to Sin3B^+/+^ LT-HSCs. **H**. GO Molecular Function was carried out for 49 genes unique to Sin3B^-/-^ LT-HSCs.

Next, we determined a pseudotime trajectory using Monocle’s graph autocorrelation analysis, utilizing a branch point in the LT-HSC cluster as an anchor (Figure 2B). We calculated a pseudotime for Sin3B^+/+^ and Sin3B^-/-^ LSKs to capture a similar path from LT-HSCs, through ST-HSCs, and finally to the various MPP subsets. Then, we calculated how gene expression changed as a function of pseudotime and grouped genes into modules that exhibited similar expression patterns. Given that the differentiation block in Sin3B^-/-^ whole bone marrow occurs at the level of LT-HSCs, we sought to document the changes in expression from LT-HSCs to ST-HSCs in our Sin3B^+/+^ dataset, and then look at perturbations in a Sin3B-dependent manner. We identified a module of genes in each genotype that displayed the highest average gene expression in LT-HSCs and decreased as cells progressed to MPPs (Figure 2C, D). We compared these modules directly to assess whether a unique transcriptional signature could justify the phenotype in LT-HSCs elicited upon Sin3B loss (Figure 2E).

We conducted Gene Ontology Analysis of the 65 genes commonly downregulated in LT-HSCs across both phenotypes and this analysis revealed an enrichment for Vitamin A metabolism (Figure 2F). This was noteworthy given the role for retinoic acid that has been recently described in the control of dormancy of LT-HSCs^32^. These data suggests that as LT-HSCs transition to ST-HSCs, both Sin3B^+/+^ and Sin3B^-/-^ cells disengage from the dormancy program.

Next, we uncovered 49 genes unique to the transition between Sin3B^+/+^ LT- and ST-HSCs. The genes uniquely expressed in Sin3B^+/+^ LT-HSCs showed an enrichment for the Wnt signaling pathway and in Rb protein regulation (Figure 2G). Wnt signaling has long been known to play a role in developmental hematopoiesis, specifically in the emergence of HSCs from the aorta-gonad-mesenephros (AGM), and in expansion of HSCs in the fetal liver, and a recent report has demonstrated a role for Wnt signaling in a dose-dependent manner to balance HSC self-renewal and differentiation^33^. The over-representation of the Rb pathway in wild-type cells is consistent with the previously published role for Sin3B in modulating Rb-E2F transcriptional repression^18,20,22^.

Finally, we analyzed the 49 genes that were uniquely expressed in Sin3B^-/-^ LT-HSCs and found enrichment for various cell-cell interaction molecules, with Cell Adhesion Molecules registering as the most significantly enriched term (Figure 2H). This observation was noteworthy given the importance of adhesion and migration in HSCs within the niche and their impact on differentiation^34^. It is tempting to speculate that the differentiation block elicited upon Sin3B inactivation stems at least in part from a defective adhesion and/or migration of LT-HSCs within the niche.

### Loss of Sin3B results in an aberrant stress response in LT-HSCs upon challenge with 5-FU

The scRNA-Seq analysis described above confirmed the subtle transcriptional deregulation of the LT-HSC compartment in Sin3B^H-/-^ mice. However, the Sin3B^-/-^ HSC phenotype only manifests during a stress response, when HSCs are actively engaging a differentiation program. To determine the Sin3B-dependent transcriptional changes in response to stress, we performed single cell transcriptome analysis on LSKs isolated from Sin3B^F/F^ and Sin3B^H-/-^ mice 9 days after 5-FU administration. 5-FU administration induces myelosuppression and subsequent activation of LT-HSCs to compensate for the elimination of rapidly proliferating progenitors that are thought to maintain homeostasis. Nine days after 5-FU administration, the number of HSCs peak before returning to homeostasis^35^, at which point HSCs are likely actively engaged in differentiation.

We applied the same computational approach to this dataset as we did for LSKs at homeostasis and recovered 4032 cells. Utilizing hashtag oligonucleotide analysis and the transcriptional signatures used for Figure 2, we assigned a genotype and hematopoietic subset identity to each cell (Figure 3A, B). When compared to mice at homeostasis, we observed a contraction in the stem cell compartment of wild-type mice exposed to stress, consistent with previous reports showing that in regenerative conditions, MPP2s and MPP3s expand to replenish myeloid output, with lymphoid regeneration taking place later^24^. When comparing across genotypes in stress conditions, we noted further expansion of the stem cell compartment in Sin3B^H-/-^ mice compared to their Sin3B^+/+^ counterparts (Figure 3C). Monocle3’s trajectory analysis revealed a more direct differentiation process in stress conditions than at homeostasis, specifically with LT-HSCs transitioning directly to MPP’s in both Sin3B^+/+^ and Sin3B^-/-^ cells (Figure 3D).

**Figure 3.**
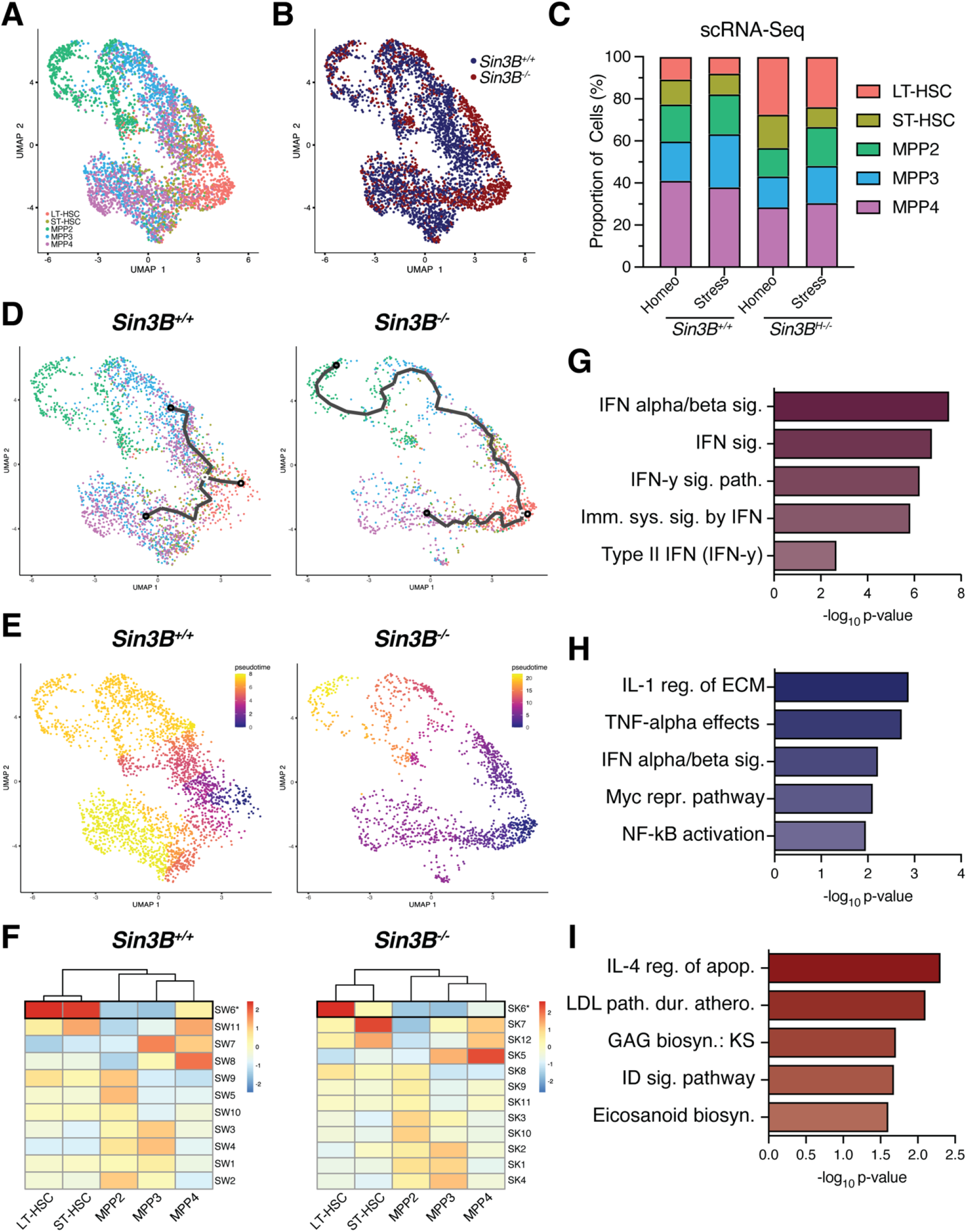
Sin3B^-/-^ LSKs display defective stress hematopoiesis. **A**. LSKs were sorted from Sin3B^F/F^ and Sin3B^H-/-^ mice 9 days after 5-FU administration (100mg/kg, i.p.) and subjected to scRNA-Seq analysis utilizing the same strategy as in Figure 1. **B**. Sin3B^+/+^ and Sin3B^-/-^ were able to be separated in our data based on hashtag oligonucleotide sequencing. **C**. Quantification of different hematopoietic subsets at indicated conditions. **D**. Monocle3 differentiation trajectory analysis of Sin3B^+/+^ and Sin3B^-/-^ LSKs during a stress response. **E**. Pseudotime analysis utilizing LT-HSCs as starting node for either Sin3B^+/+^ and Sin3B^-/-^ LSKs. **F**. Differential expression analysis was used to calculate modules of genes grouped together based on expression as a function of pseudotime. **G**. Gene Ontology analysis for 100 genes in common **H**. Gene Ontology analysis for 136 genes unique to Sin3B^+/+^ LT-HSCs. **I**. Gene Ontology analysis for 126 genes unique to Sin3B^-/-^ LT-HSCs

Next, we utilized Monocle to determine a pseudotime for each genotype along the paths previously calculated by the trajectory analysis (Figure 3E). We again reasoned that this path linking LT-HSCs to MPPs could serve as a proxy for the transcriptional landscape changes during differentiation. We then identified genes for which expression changed as a function of pseudotime in Sin3B^+/+^ LSKs upon stress (Figure 3F, left panel). The same analysis was applied to the Sin3B^-/-^ dataset (Figure 3F, right panel) resulting in the identification of two sets of genes with highest expression in LT-HSCs and decreased as cells transitioned to MPPs, either in wild-type or Sin3B^-/-^ LT-HSCs.

We identified 136 genes in the Sin3B^+/+^ data (Modules SW6) and 126 genes the Sin3B^-/-^ data (Modules SK6) (Figure 3F). The overlap between these modules revealed that 100 genes were commonly downregulated as LT-HSCs transitioned into MPPs in Sin3B^+/+^ and Sin3B^H-/-^ mice. Gene Ontology analysis indicated an enrichment for interferon signaling within these shared genes (Figure 3G). This observation is consistent with the established role for inflammation in the hematopoietic system upon stress^36^. This result suggests that both Sin3B^+/+^ and Sin3B^-/-^ LT-HSCs can discern inflammatory conditions in the bone marrow. The 36 genes unique to the wild-type LT-HSC module showed an enrichment for the NF-KB pathway (Figure 3H), known to be engaged downstream in response to hematopoietic insults^37^. Interestingly, the group of genes also displayed an enrichment for the IFN α/β signaling pathway, suggesting that while IFN signaling is engaged upon exposure to stress in Sin3B^-/-^ LT-HSCs, the intensity or the integrity of this pathway is altered in these conditions, as evidenced by the lack of TNF signaling in Sin3B^-/-^ LT-SHCs, despite expressing genes in the IFN pathway. Finally, the 26 genes unique to Sin3B^-/-^ LT-HSCs exhibited no discernable molecular function as a group (Figure 3I). Together, these data suggests that Sin3B^-/-^ LT-HSCs are likely able to sense stress conditions as a result of 5-FU treatment but are unable to engage the appropriate transcriptional differentiation programs. Of note, Sin3B^-/-^ LT-HSCs do not downregulate genes within the NF-KB pathway, which could provide a possible mechanism for their block in differentiation during a stress response.

### Sin3B restricts progression along the G_1_ phase of the cell cycle in LT-HSCs

To understand how the loss of Sin3B abrogates HSCs’ differentiation response, we first assessed the expression of master transcription factors responsible for initiation the differentiation of various lineages^38^. We found no change in expression of these transcription factors upon loss of Sin3B (Supplemental Figure 1). In light of the previously characterized Sin3B-driven repression of cell cycle genes, combined with the seemingly normal expression of components of the differentiation machinery in HSCs, we hypothesized that aberrant cell cycle progression may contribute to the inability of Sin3B^-/-^ LT-HSCs to differentiate.

To test this hypothesis, we first compared the scRNA-Seq data we obtained for Sin3B^-/-^ LT-HSCs to a recently published transcriptomic signature for quiescence^39^. Cells were ranked by progression through the cell cycle and ordered by expression levels. Loss of Sin3B resulted in a shift in the progression from quiescence towards G_1_ (Figure 4A), indicative of an impaired cell cycle control in Sin3B^-/-^ LT-HSCs. Next, we sought to test whether this spurious progression into the cell cycle bore functional consequences. We isolated and cultured LT-HSCs to assess S phase progression, by monitoring EdU incorporation at 12-hour intervals. Culture conditions for HSCs contain supraphysiological levels of cytokines and growth factors that enforce HSC cycling. Sin3B^-/-^ LT-HSCs displayed higher levels of EdU incorporation than their wild-type counterparts after 1 hour or 12 hours in culture, suggesting that Sin3B^-/-^ LT-HSCs are poised to reenter S-phase faster than wild-type cells when cultured in pro-proliferative conditions (Figure 4B). To determine if this property reflects a more advanced position within the G_0_/G_1_ phase in Sin3B^-/-^ LT-HSCs at homeostasis, we performed immunofluorescence on freshly isolated LT-HSCs. We first tested cells for the expression of p27^Kip1^, a known marker of quiescence. Loss of Sin3B resulted in lower levels of p27^Kip1^ expression in LT-HSCs, suggesting they are less quiescent than their wild-type counterparts (Figure 4C, D). As cells re-enter the cell cycle, they express Cyclin D1, which complexes with Cdk4/6 to phosphorylate the Retinoblastoma (Rb) protein and relieve repression of E2F target genes. We also detected elevated levels of Cyclin D1 in Sin3B^-/-^ LT-HSCs (Figure 4C, E). Cyclin D1 can phosphorylate Rb at multiple sites, resulting in the initiation of Rb inactivation. Indeed, immunofluorescence staining of LT-HSCs showed increased Rb (S807/811) phosphorylation in Sin3B^-/-^ LT-HSCs (Figure 4C, F). Finally, inactivation of Rb and subsequent E2F activation results in a feed-forward loop that increases the expression of Cyclin E, which complexes with Cdk2 and is a key determinant of the G_1_/S phase transition. When assaying for Cyclin E protein levels, we observed an increase in LT-HSCs upon Sin3B deletion (Figure 4C, G). In sum, these data indicate that Sin3B restricts LT-HSCs’ progression along the G_1_ phase of the cell cycle and that Sin3B^-/-^ LT-HSCs are poised to rapidly cross the G_1_/S checkpoint when stimulated to do so.

**Figure 4.**
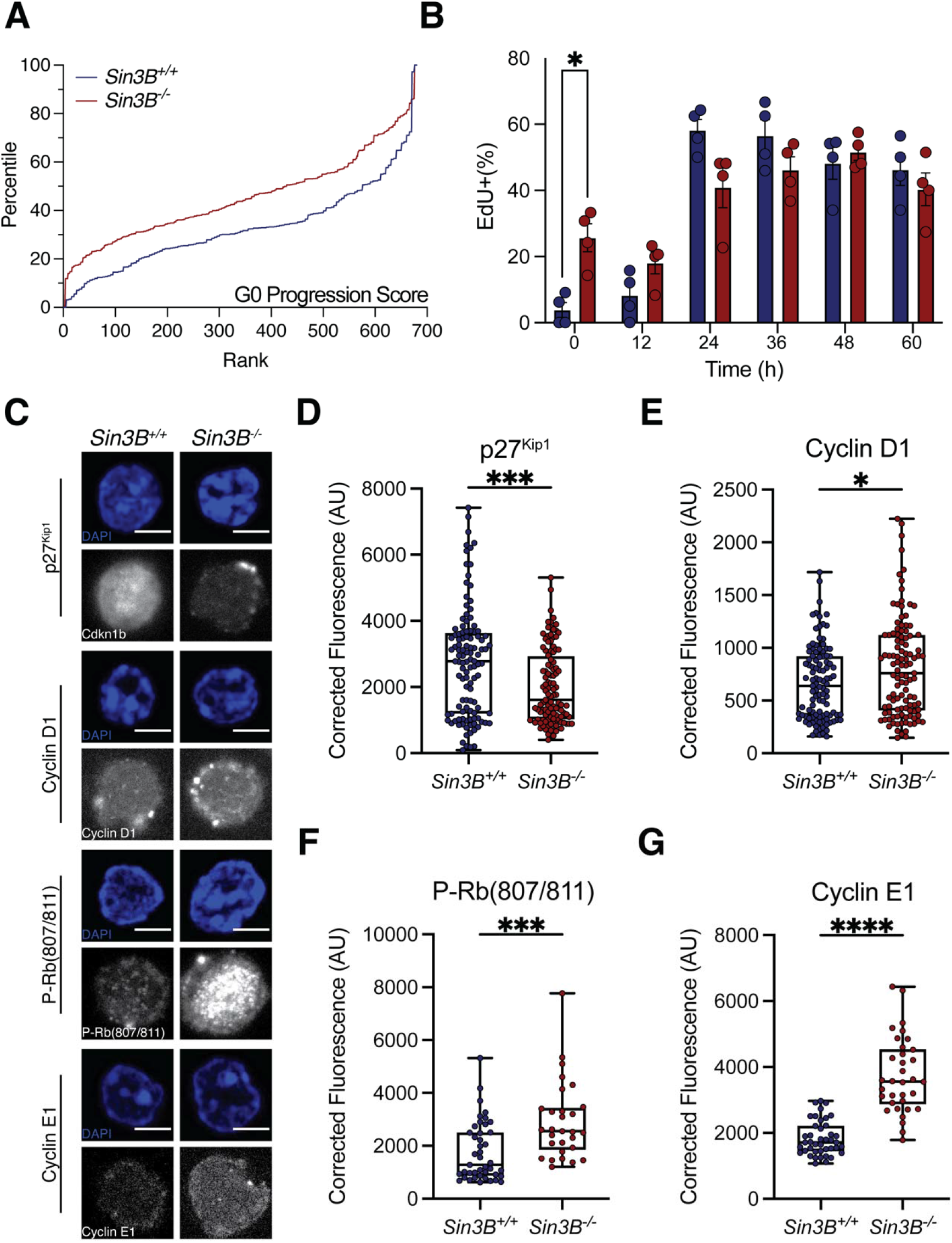
Ablation of Sin3B results in spurious cell cycle progression in LT-HSCs. **A**. G_0_ scores were calculated and cells were ranked and data transformed into percentile to normalize for cell number. **B**. EdU labeling of cycling LT-HSCs from indicated genotypes at various timepoints via immunofluorescence. Asterisks indicate statistical significance (2-way ANOVA, S □ idák’s multiple comparison’s test; *p<0.05; data shown as mean ± SEM. n = 4 per genotype. **C**. LT-HSCs were isolated from Sin3B^F/F^ or Sin3B^H-/-^ animals via FACS and processed for immunofluorescence and quantification of indicated cell cycle proteins. Signal was quantified within the nucleus of individual cells utilizing DAPI as a mask. Representative immunofluorescence from sorted LT-HSCs of indicated proteins. **D**. Quantification of Cdkn1b(p27^Kip1^) in LT-HSCs. **E**. Quantification of Phospho-Rb(S807/811) in LT-HSCs. **F**. Quantification of Cyclin D1 in LT-HSCs. **G**. Quantification of Cyclin E1 in LT-HSCs. Outliers were identified with the ROUT method (Q = 1%). Asterisks indicate statistical significance (2-tailed unpaired t-test; *p<0.05; ***p<0.0001; ****p<0.00001). n = 4 per genotype.

### Loss of Sin3B results in an aberrant chromatin environment that is restrictive to differentiation

Our central hypothesis is that aberrant cell cycle progression directly hinders the ability of LT-HSCs to differentiate. The genetic ablation of Sin3B results in de-repression of cell cycle genes and aberrant progression into the cell cycle. Sin3B^-/-^ LT-HSCs correctly express various transcription factors directing hematopoietic cell fate, and do not exhibit an overt transcriptional signature that would point to defects in differentiation at homeostasis (Supplemental Figure 1). Furthermore, after 5-FU administration, Sin3B^-/-^ LT-HSCs are capable of sensing a stress environment in the niche but remain impaired in their ability to differentiate. We therefore hypothesize that the loss of Sin3B alters the chromatin landscape that enables the transcriptional machinery to properly engage expression of the differentiation program, given that Sin3B itself is a known component of a chromatin modifying complex. To determine the impact of Sin3B on the chromatin accessibility landscape of HSCs, we performed Assay for Transposase-Accessible Chromatin (ATAC)-Seq on purified LT-HSCs from wild-type and Sin3B^-/-^ LT-HSCs at homeostasis^40^.

Our analysis identified 64,744 peaks corresponding to accessible chromatin regions in Sin3B^+/+^ LT-HSCs, and 50,655 peaks in Sin3B^-/-^ LT-HSCs. Interestingly, Sin3B^-/-^ LT-HSCs displayed overall higher levels of accessibility relative to their Sin3B^+/+^ counterparts (Supplemental Figure 2). This observation is consistent with Sin3B’s well established ability to coordinate the recruitment of histone repressors, including histone deacetylases and histone demethylases at discrete genomic loci^18,22^. Next, we utilized Diffbind in R to identify differentially accessible peaks utilizing a DESeq2 workflow with an FDR < 0.05 between the Sin3B^+/+^ and Sin3B^-/-^ LT-HSCs^41^. We uncovered 287 peaks uniquely accessible (open chromatin) in Sin3B^+/+^ LT-HSCs, compared to 5531 uniquely accessible peaks in Sin3B^-/-^ LT-HSCs (Figure 5A, B). Next, we annotated the peaks using HOMER to determine the genomic features of the loci whose accessibility is modulated by Sin3B^42^. We found no discernable differences in the respective proportion of any given genomic feature accessible in Sin3B^+/+^ and Sin3B^-/-^ LT-HSCs, such as the 5’ UTR and exons (Supplemental Figure 3A).

**Figure 5.**
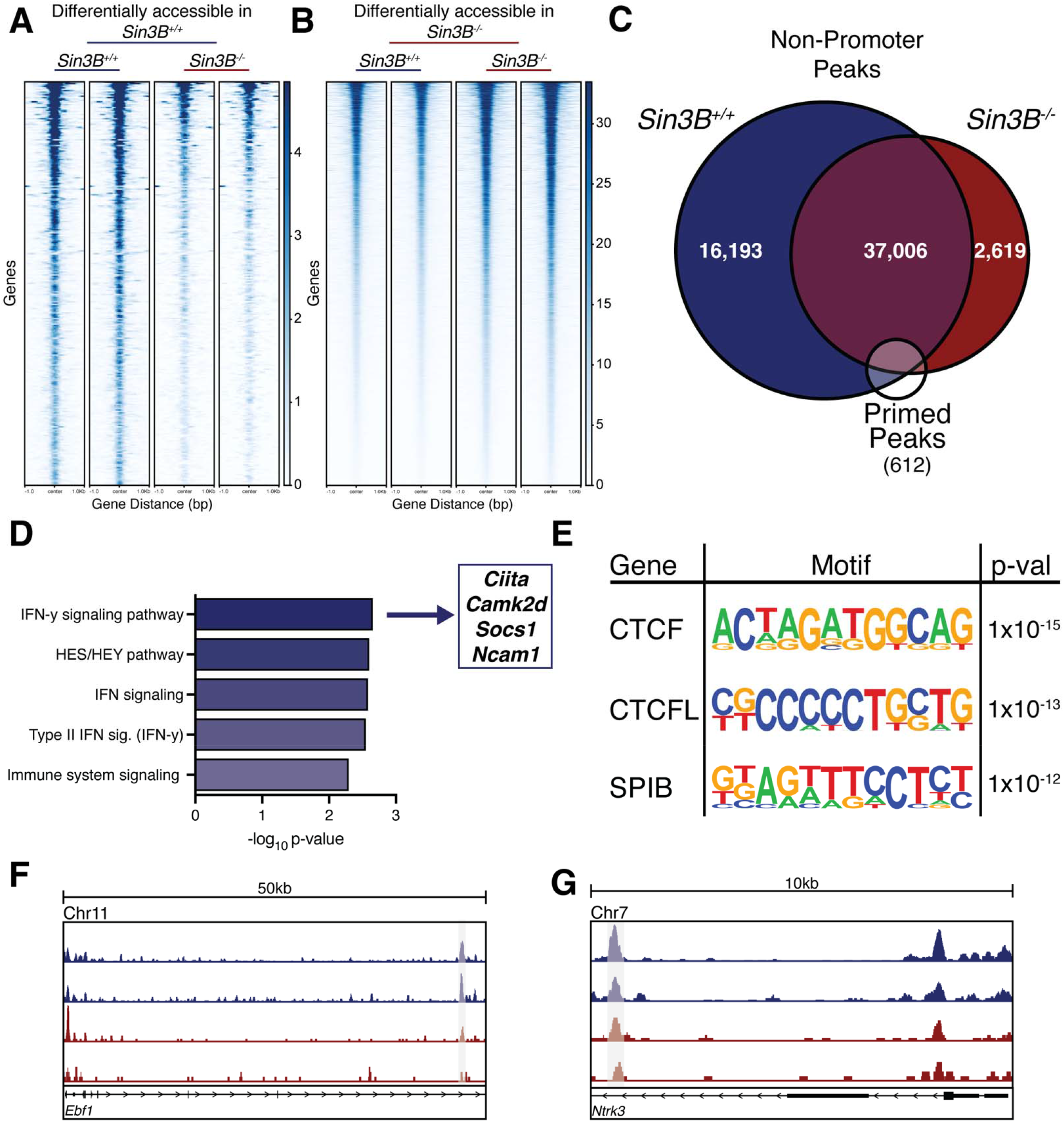
Sin3B promotes a permissive environment for differentiation. ATAC-Seq was performed on isolated LT-HSCs purified via FACS. Differentially accessible regions were calculated using the Diffbind package in R. **A**. Heatmap of differentially accessible peaks in Sin3B^F/F^ LT-HSCs. **B**. Heatmap of differentially accessible peaks in Sin3B^-/-^ LT-HSCs. **C**. Venn diagram showing overlap of non-promoter peaks from dataset, overlapped with a list of published primed peaks from Martin, et al. **D**. Primed peaks unique to wild-type LT-HSCs were analyzed with the GREAT tool and the list of genes that are putatively regulated by these regions were analyzed using Enrichr. Shown are the pathways identified by BioPlanet as significantly enriched. **E**. Homer motif analysis for primed peaks unique to Sin3B^+/+^ LT-HSCs. Shown are the top 3 motifs and p-values. **F**. Representative tracks for the gene Ebf1, a gene regulated by one of the primed peaks identified in **D** the cis-regulatory element is highlighted. **G**. Representative tracks for the gene Ntrk3, a gene regulated by one of the primed peaks identified in **D** the cis-regulatory element is highlighted.

Strikingly, when we assessed the uniquely accessible peaks in Sin3B^-/-^ LT-HSCs, the proportion of genomic features of exhibited a significant enrichment in promoter regions, at the expense of intergenic regions (Supplemental Figure 3B, C). Given that these regions are enriched for regulatory sequences, we reasoned that the differentiation defect we observe in Sin3B^-/-^ LT-HSCs may be a result of defective enhancer engagement. To test this, we used a previously published list of “primed” peaks^43^, which correspond to enhancers that are present in an active chromatin state in LT-HSCs, as well as in specific lineages of peripheral blood cells. We first filtered our data by removing all annotated promoters, and then intersected the putative regulatory regions identified as accessible in Sin3B^+/+^ or in Sin3B^-/-^ LT-HSCs with the primed peak list (Figure 5C). We recovered 493 primed peaks in Sin3B^+/+^ LT-HSCs, and 445 in Sin3B^-/-^ LT-HSCs. Next, overlapped these sets of peaks, and found 64 peaks that were unique to Sin3B^+/+^ LT-HSCs (Figure 5C, F, G). We used the GREAT tool to interrogate what genes these enhancers regulate and uncovered over 100 putatively regulated genes. Gene ontology analysis for this group of genes revealed significant enrichment for interferon gamma signaling (Figure 5D)^44^. Notably, we had detected interferon signaling as a transcriptional signature altered in the scRNA-Seq dataset in stress conditions (Figure 3H, I). Additionally, a report by Baldridge and colleagues demonstrated that interferon gamma was important in mediating LT-HSC activation and proliferation^45^. Together, these results indicate a defect in the chromatin structure of loci related to interferon signaling in Sin3B^-/-^ LT-HSCs which correlates with their to properly commit to various downstream lineages and subsequent differentiation.

Next, we used HOMER’s motif analysis to identify the putative sequence-specific transcription factors bound to the 64 primed peaks that were unique to Sin3B^+/+^ LT-HSCs to identify the putative sequence specific transcription factors bound to these sites. The top two enriched motifs corresponded to the binding sites for CTCF and the related CTCFL/BORIS (Figure 5E). These proteins have been implicated in maintaining quiescence in hematopoietic stem cells in mice, and their inactivation results in defective survival and differentiation in HSCs^46^. Additionally, Qi and colleagues demonstrated that CTCF mediates cis-regulatory element binding with their promoters as HSCs differentiate into a specific lineage, with no impact on TAD formation or boundaries^47^. Our results indicate that the accessibility to CTCF binding sites that control HSC differentiation is impaired in the absence of Sin3B. As previous studies have demonstrated that dynamic CTCF binding is required for lineage specific differentiation in human HSCs, to mediate the transition into “activation” in human HSPCs, it is tempting to speculate that the differentiation of Sin3B^-/-^ HSCs is functionally linked to the compromised accessibility of these CTCF binding sites. Together, our data suggest that Sin3B modulates the chromatin state of CTCF-bound sites that are important for lineage priming in LT-HSCs.

## Discussion

HSCs must balance differentiation capabilities with self-renewal and proliferation potential to meet the need of an organism throughout its lifespan. To maintain these stemness features, HSCs normally reside in a quiescent state, a temporary form of cell cycle exit, termed G_0_. Proliferation and self-renewal require re-entry into the cell cycle, making early cell cycle progression an intrinsic mechanism of HSC function. We demonstrate here that the chromatin-associated Sin3B protein potentiates differentiation in HSCs, correlating with its ability to restrict progression within the G_1_ phase of the cell cycle. This is supported by data showing a spurious progression into G_1_ cells in HSCs genetically inactivated for Sin3B, driven by the expression of canonical G_1_ progression mediators such as Cyclin E2. Additionally, scRNA-Seq analysis reveals that Sin3B^-/-^ LT-HSCs appropriately express transcription factors required for lineage commitment at homeostasis, or under stress conditions, but are unable to engage the corresponding pro-differentiation response. Our evidence suggests that Sin3B fine tunes the control mechanisms of HSC differentiation, and its loss results in an inability of the cell to regulate differentiation under stress.

It has been previously demonstrated that mouse and human embryonic stem cells exhibit different levels of response to differentiation stimuli depending on the phase of the cell cycle they reside in^11^, raising the question: How does progression through cell cycle mechanistically potentiate differentiation potential? Data from ES cells have demonstrated that after mitosis, specific regulatory sequences driving self-renewal and stemness contact their target genes earlier in G_1_, unlike regulatory sequences controlling differentiation^48^. These observations suggest a prioritization of chromatin unfolding whereby the spatial organization of genes related to cellular identity is quickly re-established before other transcriptional programs can be engaged. Once cells commit to proliferation and initiate S phase, the chromatin landscape is restructured to prepare for genome replication. Here, we postulate the existence of a “commitment window” that begins when chromatin unfolds after mitosis and ends when chromatin is reconfigured for DNA replication. Our data supports that this mechanism might be responsible for potentiating differentiation in adult stem cells.

This hypothesis is supported by data showing that overexpression of CCND1-CDK4 in human HSCs confers a competitive advantage *in vivo*, as they transition from G_0_ to G_1_, and are refractory to differentiation signals *in vitro*. The authors demonstrated that expression of the CCND1-CDK4 construct resulted in a shorter G_1_ phase correlating with enhanced self-renewal^14^. Additional evidence comes from a study demonstrating that the speed through which murine Megakaryocytic-erythroid progenitors (MEPs) transit the cell cycle mediate the bifurcation towards either Megakaryocytic Progenitors (MkPs) or Erythroid Progenitors (ERPs)^16^. Additionally, our own data demonstrates that as a result of spurious cell cycle progression, Sin3B^-/-^ LT-HSCs display aberrant transcriptional programs related to differentiation, such as their inability to downregulate the Wnt pathway as they transition to ST-HSCs, and their improper expression of cell adhesion molecules. Of note, Sin3B^-/-^ LT-HSCs appear to improperly express and regulate genes in the interferon pathway, which have been established to play a crucial role in HSC activation and differentiation. Of note, Sin3B^-/-^ LT-HSCs appear to improperly express and regulate some genes in the interferon pathway (Figure 3G), which is consistent with an impaired chromatin accessibility profile at these loci (Figure 5D), which have been established to play a crucial role in HSC activation and differentiation.

Interferons, a group of inflammatory cytokines, have been established to play an important role in HSC biology, both during development and at homeostasis or during stress responses^49^. Both class I and II interferons have been found to directly signal to HSCs, and conflicting reports have emerged categorizing these cytokines as either promoting or suppressing hematopoiesis, depending on the experimental design. However, recent studies utilizing direct treatment of mice with IFNα^50^, or IFNγ^51^ and analysis of HSCs *in vivo* have shown that these cytokines induce cell cycle entry of quiescence HSCs. Of note, such treatments with these cytokines stimulate an exit from quiescence, but the overall HSC pool does not increase in number, as a result of impaired self-renewal^52^. Additionally, treatment with IFNα and IFNγ can promote megakaryocytic^53^ and myeloid differentiation^54^, respectively. Together, these observations could at least in part justify the impaired differentiation elicited by Sin3B inactivation in LT-HSCs. Of note, the relationship between cell cycle position and IFN signaling status remains to be investigated in HSCs.

Mechanistically, a report in human HSCs demonstrated the transition from LT- to ST-HSCs was associated with cell cycle re-entry and changes in CTCF binding sites. These changes altered 3D chromatin interactions in repress stemness genes in ST-HSCS normally expressed in LT-HSCs^8^. In light of the recent development of small molecule inhibitors of early cell cycle progression in clinical settings, these studies point to cell cycle progression as actionable opportunity for the modulation of hematopoietic stem and progenitor cells expansion/differentiation decision.

## Acknowledgements

We thank all members of the David laboratory for their feedback throughout this project, the NYU Genome Technology Center (RRID: SCR_017929) for expertise with sequencing experiments, and the Cytometry and Cell Sorting Laboratory (RRID: SCR_019179) for providing the flow cytometry technologies, and the High Performance Computing Core for computational requirements. We thank Camilla Forsberg and Eric Martin for the primed peak list; Eftychia Apostolou, Markus Schober, William L. Carroll, and Christopher Y. Park for helpful discussions. The cores are supported in part by National Institutes of Health (NIH) Grant P30CA016087. A.C. is supported by the National Cancer Institute (NCI) (F31CA232659). G.D. is supported by the NCI/NHLBI (R01CA148639, R21CA206013, R21CA246416, R56HL163940), the New York State Department of Health (C36617GG).

## Author Contributions

Conceptualization, A.C. and G.D.; Methodology, A.C., F.B., K.V.R, and G.D.; Formal Analysis, A.C., T.M., K.V.R., and G.D.; Investigation, A.C., T.M, and F.B.; Resources, K.V.M and G.D.; Writing – Original Draft, A.C. and G.D.; Writing – Review & Editing, A.C. and G.D.; Visualization, A.C., Supervision, K.V.R. and G.D.

## Inclusion and Diversity Statement

One or more of the authors of this paper self-identifies as an underrepresented ethnic minority in science. One or more of the authors of this paper self-identifies as a member of the LGBTQ+ community. One or more of the authors of this paper received support from a program designed to increase minority representation in science.

## Materials and Methods

### Mice

Mice containing the Sin3B-flox (*Sin3B*^*F*^) allele have been previously described. To generate hematopoietic specific deletion of Sin3B, *Sin3B*^*F/F*^ mice were intercrossed to *Vav1-iCre* mice, which is active at embryonic day 11.5 (E11.5). *Ptprc*^*a*^; *Pepc*^*b*^ (CD45.1) congenic mice were purchased from the Jackson Laboratory and bred in house to use as recipients for competitive transplantation experiments. All mice were kept on an inbred C57BL/6 background. Mice were housed in pathogen-free barrier facilities with a 12-hour light/dark cycle and given food and water *ad libitum*. Mice were administered 5-Fluorouracil (Invivogen) via intraperitoneal injection at 100mg/kg body weight. For competitive transplantation assays, recipient mice at 8-10 weeks of age were lethally irradiated (total body irradiation) with 12Gy of γ-irradiation with a MultiRad 350 X-Ray Irradiator (Faxitron®). Mice were given 2 doses of 6Gy of irradiation at least 3 hours apart. Mice were maintained on sterile, acidified water supplemented with Sulfamethoxazole and Trimethoprim for two weeks following irradiation, replenishing the antibiotics after a week. Equal numbers of male and female mice were used in all experiments unless specified otherwise. All animal experiments and protocols were approved by the New York University Grossman School of Medicine Institutional Animal Care and Use Committee.

### Flow cytometry and cell sorting

To isolate indicated cell populations, mice were humanely sacrificed via CO_2_ inhalation and cervical dislocation was used as a secondary means of euthanasia. Femurs, tibiae, and pelvis were isolated from mice, and if increased numbers of cell were required, sternum, humeri, and vertebrae was dissected as well. Whole bone marrow was isolated from bones (femurs, tibiae, pelvis) through spinning in a microcentrifuge for 8 seconds into FACS-E buffer (1X phosphate buffered saline [PBS] supplemented with 2% fetal bovine serum [FBS] and 25mM ethylenediaminetetraacetic acid [EDTA]) or through crushing in a mortar and pestle (sternum, humeri, vertebrae).

For whole bone marrow analysis, whole bone marrow was incubated in Ammonium-Chloride-Potassium (ACK) lysis buffer for 5 minutes on ice to remove erythrocytes. Cells were then resuspended in FACS buffer (1X PBS supplemented with 2% FBS) and incubated with a cocktail of biotinylated antibodies against lineage markers and Rat IgG (20μg/mL) for 30 minutes on ice. Cells are washed and then incubated with HSPC antibodies conjugated to fluorophores and Rat IgG for 90 minutes. Cells are washed and resuspended in FACS buffer carrying 4’,6-diamidino-2-phenylindole (DAPI, 500ng/mL) to mark dead cells and analyzed on either a Bectin, Dickinson, and Company (BD™) LSR II UV (equipped with 355nm, 407 nm, 488nm, 561nm, 633nm lasers) or a BD™ LSR II HTS (equipped with 407nm, 488nm, 561nm, 633nm lasers). Data collection was done using BD FACSDiva™ software and .fcs files were formally analyzed with FlowJo (FlowJo, BD). All flow cytometry experiments contained single color controls for compensation and gating.

For fluorescence activated cell sorting, whole bone marrow was first blocked with TruStain FcX™ PLUS (anti-mouse CD16/32) (50μg/mL) for 5 minutes on ice in MACS Buffer (1X PBS supplemented with 1% FBS, 1% bovine serum albumin [BSA], and 2mM EDTA that was sterile filtered through 0.22μm filter and de-gassed). Then, cells were incubated with anti-mouse CD117 microbeads (20μL for femurs, tibiae, and pelvis, 40μL if also isolating cells from sternum, humeri, and vertebrae) for 15 minutes on ice. Cells were washed in MACS buffer and then filtered through a 40μm mesh before being loaded onto a Miltenyi Biotec MS column placed in a miniMACS separator. Flowthrough containing CD117^-^ cells was discarded. MS column was washed with MACS buffer and then flowthrough discarded. Column was then removed from magnet and placed in a microcentrifuge tube. MACS buffer was loaded and plunger used to gently expel cells from the column. This CD117^+^ enriched fraction was then stained as previously described for lineage markers and HSPC markers before being resuspended in FACS buffer containing DAPI and passed through a 40μm filter again and sorted on a BD FACSAria™ II (equipped with 355nm, 407nm, 488nm, 561nm, 633nm lasers) or a BD FACSAria™ IIu SORP (equipped with 355nm, 407nm, 488nm, 561nm, 633nm lasers) utilizing a 100μm nozzle. Single color controls were used for compensation and florescence minus one controls were utilized to set gates before sorting using FACSDiva software. Cells were sorted into 1X PBS supplemented with 2% FBS. Analysis of sorting data was accomplished with FloJo software. Visualization and statistical analysis was computed after exporting data to GraphPad Prism9 software.

### Hematopoietic Stem and Progenitor Immunophenotypes

The following markers were used for the indicated cell types:

LT-HSC: L^-^S^+^K^+^Flk2^-^CD48^-^CD150^+^

ST-HSC: L^-^S^+^K^+^Flk2^-^CD48^-^CD150^-^

MPP2: L^-^S^+^K^+^Flk2^-^CD48^+^CD150^+^

MPP3: L^-^S^+^K^+^Flk2^-^CD48^+^CD150^-^

MPP4: L^-^S^+^K^+^Flk2^+^CD48^+^CD150^-^

L (Lineage) markers: CD3, CD4, CD8a, CD11b, B220, Gr1, IL-7Rα, Ter-119 S: Sca-1 (Ly6a)

K: c-Kit (CD117)

Flk2: Flt-3 (CD135)

### Competitive transplantation assay

Recipient CD45.1 mice at 8-10 weeks of age were irradiated the day before transplantation experiments, with split doses of irradiation at least 3 hours apart. Donor *Sin3B*^*F/F*^ or *Sin3B*^*H-/-*^ were used at 6-8 weeks of age. Whole bone marrow was isolated erythrocytes lysed as described in Flow Cytometry and analysis section. Whole bone marrow cells were counted manually using a hemacytometer. 1×10^6^ donor wild-type or *Sin3B*^*-/-*^ cells were mixed at a 1:1 ratio with wild-type competitor CD45.1. Cells were washed with 1X PBS to remove traces of serum and 2×10^6^ cells were resuspended in 100μL sterile 0.22μm filtered 1X PBS and transplanted into mice via retroorbital injection using a 31G, 6mm insulin syringe (BD). After 8 weeks, mice were sacrificed and whole bone marrow was isolated and stained as described above for HSPC markers with the addition of antibodies to distinguish between CD45.1 and CD45.2 alleles. Data was analyzed using FloJo and statistically analyzed in Prism9.

### EdU incorporation assay

LT-HSCs were sorted as described above, and cultured in 96 well round bottom plates containing 100μL of HSPC media (5% FBS, Stem Cell Factor [SCF, 25ng/mL], Interleukin-11 [IL-11, 25ng/mL], FMS-like tyrosine kinase 3 ligand [Flt-3L, 25ng/mL], Thrombopoietin [TPO, 25ng/mL], Interleukin-3 [IL-3, 10ng/mL], Granulocyte-macrophage colony-stimulating factor [GM-CSF, 10ng/mL], Erythropoietin [EPO, 4Units/mL], 1% penicillin G/streptomycin, 2% GlutaMAXTM, 55μM 2-mercaptoethanol in Iscove’s Modified Dulbecco’s Medium [IMDM]) for indicated time periods in a 37ºC humidified water-jacketed cell culture incubator with 5% CO_2_. At timepoints, cells were given 100μL of fresh HSPC media containing 20μM 5-Ethynyl-2’-deoxyuridine (EdU) for a final concentration of 10μM. Cells were incubated for one hour, and then plated onto poly-L-lysine coated #1.5 coverslips placed in individual wells of 12 well plates. Cells were allowed to attach for 15 minutes at room temperature (RT). Then 800μL of BD Cytofix™ buffer was added to fix cells for 10 minutes at RT with gentle agitation. Then 200μL of 1M Glycine in ddH_2_O was added to quench fixation. Cells were washed three times with 1X PBS before proceeding to EdU staining.

Cells were processed with the Click-iT® Plus EdU Imaging Kit. After washing, cells were permeabilized with Triton X-100 Buffer (0.5%[v/v] Triton X-100; 20mM HEPES-KOH, pH 7.9; 50mM NaCl; 3mM MgCl_2_; 300mM sucrose; 0.05%[w/v] NaN_3_ in ddH2O) for 10 minutes at RT with gentle agitation. Cells were washed twice with IF Washing Buffer (1% FBS; 1% BSA; 0.1% Triton X-100; 0.1%[v/v] Tween-20; 0.05% NaN_3_ in 1X PBS) before being incubated with Click-iT® reaction cocktail containing Alexa Fluor® 488 picolyl azide. Cells were incubated with reaction cocktail for 30min at RT protected from light. Samples were then washed with IF Washing buffer containing 500ng/mL DAPI, and then washed 3 more times with IF Washing buffer, and then once with 1X PBS, before being mounted on slides with Vectashield® (Vector Labs). Slides were sealed with commercially available clear nail polish and allowed to dry before being imaging on an inverted Zeiss LSM 700 Laser Scanning Confocal Microscope (equipped with 405nm, 488nm, 555nm, 639 lasers) using a 63X plan apochromat 1.4 oil objective. Zen software was used to acquire images, using 3.0 zoom and preventing saturation of images. Images were exported to FIJI (Fiji is just ImageJ) to quantify proportion of cells staining positively for EdU. Quantification was exported to Prism9 for additional analysis and visualization.

### Immunofluorescence of LT-HSCs

LT-HSCs from *Sin3B*^*F/F*^ and *Sin3B*^*H-/-*^ mice were isolated via FACS and plated on poly-L-lysine coverslips as described above for EdU labeling. After permeabilization, cells were washed twice with 1X PBS, and then blocked with 1X PBS supplemented with 5% (v/v) goat serum and 0.1% (v/v) Tween-20 for one hour at RT with gentle agitation. Coverslips were then flipped onto 100μL droplets of blocking buffer (10% FBS, 2.5% BSA, 0.1% Tween-20, 0.1% Triton X-100, 0.05% NaN_3_ in 1X PBS) containing primary antibody at a 1:100 dilution. Cells were incubated with primary antibody for one hour at RT. Coverslips were then returned to individual wells of 12 well plate and washed 3 times with IF washing buffer. Coverslips were flipped again onto droplets of blocking buffer containing secondary antibody. Goat anti-rabbit Alexa Fluor 488 or Goat anti-mouse Alexa Fluor 594 antibodies (Invitrogen) at a 1:400 dilution were used and cells were incubated for 1 hour at RT protected from light. Cells were then washed 3 times in IF buffer, with the first wash carrying DAPI (500ng/mL). Coverslips were washed once with 1X PBS before being mounted on slides in Vectashield. Samples were sealed with nail polish and imaged using a Zeiss 700 as detailed above. Z-stacks were taken and the widest slice of each cell was utilized for quantification using FIJI. A mask was manually drawn using DAPI to quantify the signal for the indicated antibody within the nucleus. A small area not containing cells was quantified to determine background. Corrected fluorescence was determined by the following formula: CorrFluor = Sample – (Area x Background). All data was exported to Prism9 for statistical analysis.

### Single cell RNA-Sequencing: cDNA library preparation and sequencing

For homeostasis dataset, whole bone marrow was isolated as described above, and antibodies against Lineage, Sca1, and cKit were for 2 *Sin3B*^*F/F*^ and 2 Sin3B^H-/-^ male mice. LSKs were pooled and cells were blocked again with TruStain FcX PLUS for 5 minutes before wild-type cells were incubated with TotalSeq™-B0096 anti-mouse CD45 antibody and knockout cells with TotalSeq™-B0157 anti-mouse CD45.2 antibody for 15 minutes. Cells were washed with FACS buffer before being manually counted with a hemacytometer. The 10X Genomics Chromium Single Cell 3’ v3 kit was used to generate single cell suspensions. After counting, 2.5×10^4^ cells from each genotype were combined with MasterMix and loaded onto a Chromium Chip along with the gel beads and partitioning oil. Gasket was carefully placed over chip and was loaded into a Single Cell Controller. GEMs were carefully pipetted out and visually inspected before being placed in a thermal cycler for cDNA synthesis with the following parameters (Step 1 – 53ºC for 45 minutes; Step 2 – 85ºC for 5 minutes; Step 3 – 4ºC: Hold)

Reaction was stored at -20ºC until cDNA library preparation. Quality of cDNA was checked with an Agilent 2100 Bioanalyzer. Libraries were generated via using the 10X Genomics 3’ GEM protocol with HTO primers and sequenced on an Illumina NovaSeq.

For dataset of LSKs at stress, the same workflow was utilized, except mice were first given a single dose of 5-fluorauracil (100mg/kg) injected intraperitoneally. After 9 days, mice were sacrificed and bone marrow was processed and stained as previously described and LSKs were isolated via FACS utilizing the same strategy.

#### Single cell RNA-Seq: Analysis

After sequencing, reads from cDNA and Hashtag oligos (HTOs) were demultiplexed and aligned using the 10X Genomics CellRanger software. This generated a matrix file, features file, and barcodes file that was imported into R. The Seurat package was used for downstream analysis. Briefly, the output files from CellRanger were used to generate a Seurat object. HTO counts were extracted and added to the metadata of captured cells. Data was normalized and the HTODemux function was used to classify cells. Only singlets were kept for downstream analysis. Quality control was used to filter to calculate the distribution of genes per cell identified, as well as overall counts and the proportion of reads coming from the mitochondria. Cells in the top and bottom 2% of these metrics were filtered out. Cells found to be expressing any lineage markers used in the FACS isolation were also removed. Additionally, cells expressing genes related to biased and lineage-primed HSCs were also removed.

Next, LSKs were assigned a subset identify utilizing previously published transcriptional signatures of LT-HSCs, ST-HSCs, MPP2s, MPP3s, and MPP4s. To generate Uniform Manifold Approximation Projections (UMAPs) data was scaled and Principal Component Analysis (PCA) was computed. JackStraw was then used to calculate statistically significant PC’s to use for UMAP analysis. Differentially expressed genes were determined using the FindMarkers function. The dataset containing LSKs at stress was analyzed using the same workflow as just described.

To integrate the homeostasis and stress datasets, we utilized Seurat’s IntegrateData function. First, variable features were calculated for both datasets, and integration anchors were calculated using FindIntegrationAnchors. These anchors were then used for the IntegrateData function to generate the object containing cells from homeostasis and stress.

For pseudotime analysis, the Monocle3 package was used. First, the relevant Seurat object was converted into a CellDataSet format using the as.cell_data_set function from the SeuratWrappers package. The UMAP calculation and LSK subsets defined in Seurat were used in the learn_graph function of Monocle3 when determining cell trajectories. The order_cells function was used to determine pseudotime, with the node containing the most LT-HSCs manually selected as the beginning of the pseudotime trajectory. The graph autocorrelation analysis was completed using the graph_test function using the “principal_graph” that constituted the previously calculated trajectory in learn_graph and order_cells. Genes with a q_value < 0.05 were selected and modules of co-regulated genes as a function of pseudotime were determined via find_gene_modules. Aggregate expression of genes within individual modules was accomplished via the aggregate_gene_expression function and graphed. Modules were exported and gene lists uploaded to Enrichr in order to determine gene ontologies.

For cell cycle analysis, cell cycle scores utilizing previously published datasets were calculated using log10 transformed expression. LT-HSCs were ordered from M/G_1_, G_0_, G_0_/G_1_, G_1_/S, S, G_2_/M, and M. Cells were ordered from lowest to highest expression of their G_0_ score, and data were transformed into percentile ranks to normalize for cell number.

### Assay for Transposase-Accessible Chromatin using sequencing (ATAC-Seq)

To conduct ATAC-Seq on LT-HSCs, we utilized the ATAC-Seq kit from ActiveMotif with the following modifications. LT-HSCs from 2 mice per genotype in duplicate were pooled after sorting via FACS. The amount of Assembled Transposomes was scaled based on the number of cells we were able to isolate. After tagmentation for 30 minutes, the rest of the kit was followed per manufacturer’s instructions. Briefly, DNA Purification Binding Buffer was added to samples and transferred to a DNA binding column. Columns were washed and DNA was eluted for subsequent PCR amplification. Illumina’s indexed i7 and i5 Nextera primers were used to distinguish between samples. DNA was amplified using Q5 polymerase and specific primer combinations for 10 cycles with the following conditions on a thermal cycler (Step 1 – 72ºC, 5 minutes; Step 2 – 98ºC, 30 seconds; Step 3 – 98ºC, 10 seconds; Step 4 – 63ºC, 30 seconds; Step 5 – 72ºC, 1 min; Repeat Steps 3 – 5 nine more times for a total of 10 cycles; Step 6 – 10ºC, hold). SPRI (Solid Phase Reversible Immobilization) beads were used for clean-up. Beads were washed twice with ethanol and DNA was eluted from beads. Size distribution of libraries was determined using a TapeStation, and concentration with Bioanalyzer. Our samples required additional PCR cycles and the same process was repeated on libraries for an additional 2 cycles before bead clean-up was repeated. Libraries were sequenced on an Illumina NovaSeq. Fastq files were run through FastQC and trimmed. Reads were then aligned using bowtie2 and duplicates were removed using sambamba. Bigwig files were generated using deeptools and peaks were called using MACS2.

Bed files containing peaks were imported into R and the DiffBind package was used for downstream analysis. Peaksets were read in and normalized before differential analysis was calculated using DESeq2. The HOMER package was used to annotate peaks and to calculate enrichment of DNA-binding factor motifs. The bedtools suite was utilized to compare peaksets to each other, with the intersect function used to directly compare lists of accessible peaks. Peaks were then fed into the GREAT tool using default parameters to determine putative genes regulated by the accessible chromatin peaks we identified. Finally, those genes were then used as input for Enrichr to determine Gene Ontology enrichment. Individual ATAC-Seq tracks were loaded by opening bigwig files in IGV. For motif analysis, the HOMER function findMotifsGenome.pl was used for individual peak lists. Primed peaks list were taken from Martin, et al.^43^ and first changed to an mm10 annotation format using the UCSC genome browser tool. Lists were directly compared to ATAC-seq peaks using bedtools.

### Statistical Considerations

Samples were compared using the statistical test indicated in figure legend.

Sample sizes were not determined with any formal power calculation.

## Bibliography

1. Orkin, S. H. & Zon, L. I. Hematopoiesis: An Evolving Paradigm for Stem Cell Biology. Cell 132, 631–644 (2008).

2. Haas, S., Trumpp, A. & Milsom, M. D. Causes and Consequences of Hematopoietic Stem Cell Heterogeneity. Cell Stem Cell 22, 627–638 (2018).

3. Morrison, S. J. & Scadden, D. T. The bone marrow niche for haematopoietic stem cells. Nature 2014 505:7483 505, 327–334 (2014).

4. Nakamura-Ishizu, A., Takizawa, H. & Suda, T. The analysis, roles and regulation of quiescence in hematopoietic stem cells. Development 141, 4656–4666 (2014).

5. Passegué, E., Wagers, A. J., Giuriato, S., Anderson, W. C. & Weissman, I. L. Global analysis of proliferation and cell cycle gene expression in the regulation of hematopoietic stem and progenitor cell fates. Journal of Experimental Medicine 202, 1599–1611 (2005).

6. Wilson, A. et al. Hematopoietic Stem Cells Reversibly Switch from Dormancy to Self-Renewal during Homeostasis and Repair. Cell 135, 1118–1129 (2008).

7. Laurenti, E. et al. CDK6 Levels Regulate Quiescence Exit in Human Hematopoietic Stem Cells. Cell Stem Cell 16, 302–313 (2015).

8. Takayama, N. et al. The Transition from Quiescent to Activated States in Human Hematopoietic Stem Cells Is Governed by Dynamic 3D Genome Reorganization. Cell Stem Cell 28, 488–501.e10 (2021).

9. Soufi, A. & Dalton, S. Cycling through developmental decisions: how cell cycle dynamics control pluripotency, differentiation and reprogramming. Development 143, 4301–4311 (2016).

10. Savatier, P., Huang, S., Szekely, L., Wiman, K. G. & Samarut, J. Contrasting patterns of retinoblastoma protein expression in mouse embryonic stem cells and embryonic fibroblasts. Oncogene 9, 809–18 (1994).

11. Coronado, D. et al. A short G1 phase is an intrinsic determinant of naïve embryonic stem cell pluripotency. Stem Cell Res 10, 118–131 (2013).

12. Koledova, Z. et al. Cdk2 Inhibition Prolongs G1 Phase Progression in Mouse Embryonic Stem Cells. Stem Cells Dev 19, 181–194 (2010).

13. Calder, A. et al. Lengthened G1 Phase Indicates Differentiation Status in Human Embryonic Stem Cells. Stem Cells Dev 22, 279–295 (2013).

14. Mende, N. et al. CCND1–CDK4–mediated cell cycle progression provides a competitive advantage for human hematopoietic stem cells in vivo. Journal of Experimental Medicine 212, 1171–1183 (2015).

15. Grinenko, T. et al. Hematopoietic stem cells can differentiate into restricted myeloid progenitors before cell division in mice. doi:10.1038/s41467-018-04188-7.

16. Lu, Y.-C. et al. The Molecular Signature of Megakaryocyte-Erythroid Progenitors Reveals a Role for the Cell Cycle in Fate Specification. Cell Rep 25, 2083–2093.e4 (2018).

17. Cantor, D. J. & David, G. The chromatin-associated Sin3B protein is required for hematopoietic stem cell functions in mice. Blood 129, 60–70 (2017).

18. David, G. et al. Specific requirement of the chromatin modifier mSin3B in cell cycle exit and cellular differentiation. Proceedings of the National Academy of Sciences 105, 4168–4172 (2008).

19. Rielland, M. et al. Senescence-associated SIN3B promotes inflammation and pancreatic cancer progression. J Clin Invest 124, 2125–2135 (2014).

20. Grandinetti, K. B. et al. Sin3B Expression Is Required for Cellular Senescence and Is Up-regulated upon Oncogenic Stress. Cancer Res 69, 6430–6437 (2009).

21. DiMauro, T., Cantor, D. J., Bainor, A. J. & David, G. Transcriptional repression of Sin3B by Bmi-1 prevents cellular senescence and is relieved by oncogene activation. Oncogene 34, 4011–4017 (2015).

22. Bainor, A. J. et al. The HDAC-Associated Sin3B Protein Represses DREAM Complex Targets and Cooperates with APC/C to Promote Quiescence. Cell Rep 25, 2797–2807.e8 (2018).

23. Oguro, H., Ding, L. & Morrison, S. J. SLAM Family Markers Resolve Functionally Distinct Subpopulations of Hematopoietic Stem Cells and Multipotent Progenitors. Cell Stem Cell 13, 102–116 (2013).

24. Pietras, E. M. et al. Functionally Distinct Subsets of Lineage-Biased Multipotent Progenitors Control Blood Production in Normal and Regenerative Conditions. Cell Stem Cell 17, 35–46 (2015).

25. Stoeckius, M. et al. Cell Hashing with barcoded antibodies enables multiplexing and doublet detection for single cell genomics. Genome Biol 19, 224 (2018).

26. Pei, W. et al. Resolving Fates and Single-Cell Transcriptomes of Hematopoietic Stem Cell Clones by PolyloxExpress Barcoding. Cell Stem Cell 27, 383–395.e8 (2020).

27. Polak, R. & Buitenhuis, M. The PI3K/PKB signaling module as key regulator of hematopoiesis: implications for therapeutic strategies in leukemia. Blood 119, 911–923 (2012).

28. Cancelas, J. A. & Williams, D. A. Rho GTPases in hematopoietic stem cell functions. Curr Opin Hematol 16, 249–254 (2009).

29. Florian, M. C. et al. Aging alters the epigenetic asymmetry of HSC division. PLoS Biol 16, e2003389 (2018).

30. Trapnell, C. et al. The dynamics and regulators of cell fate decisions are revealed by pseudotemporal ordering of single cells. Nat Biotechnol 32, 381–386 (2014).

31. Macaulay, I. C. et al. Single-Cell RNA-Sequencing Reveals a Continuous Spectrum of Differentiation in Hematopoietic Cells. Cell Rep 14, 966–977 (2016).

32. Cabezas-Wallscheid, N. et al. Vitamin A-Retinoic Acid Signaling Regulates Hematopoietic Stem Cell Dormancy. Cell 169, 807–823.e19 (2017).

33. Luis, T. C. et al. Canonical Wnt Signaling Regulates Hematopoiesis in a Dosage-Dependent Fashion. Cell Stem Cell 9, 345–356 (2011).

34. Zhang, J. et al. In situ mapping identifies distinct vascular niches for myelopoiesis. Nature 590, 457–462 (2021).

35. Baumgartner, C. et al. An ERK-Dependent Feedback Mechanism Prevents Hematopoietic Stem Cell Exhaustion. Cell Stem Cell 22, 879–892.e6 (2018).

36. Takizawa, H., Boettcher, S. & Manz, M. G. Demand-adapted regulation of early hematopoiesis in infection and inflammation. Blood 119, 2991–3002 (2012).

37. Bottero, V., Withoff, S. & Verma, I. M. NF-κB and the regulation of hematopoiesis. Cell Death & Differentiation 2006 13:5 13, 785–797 (2006).

38. Cabezas-Wallscheid, N. et al. Identification of Regulatory Networks in HSCs and Their Immediate Progeny via Integrated Proteome, Transcriptome, and DNA Methylome Analysis. Cell Stem Cell 15, 507– 522 (2014).

39. Chang, C. A. et al. Ontogeny and Vulnerabilities of Drug-Tolerant Persisters in HER2+ Breast Cancer. Cancer Discov 12, 1022–1045 (2022).

40. Corces, M. R. et al. An improved ATAC-seq protocol reduces background and enables interrogation of frozen tissues. Nat Methods 14, 959–962 (2017).

41. Ross-Innes, C. S. et al. Differential oestrogen receptor binding is associated with clinical outcome in breast cancer. Nature 481, 389–393 (2012).

42. Heinz, S. et al. Simple Combinations of Lineage-Determining Transcription Factors Prime cis-Regulatory Elements Required for Macrophage and B Cell Identities. Mol Cell 38, 576–589 (2010).

43. Martin, E. W. et al. Chromatin accessibility maps provide evidence of multilineage gene priming in hematopoietic stem cells. Epigenetics Chromatin 14, 1–15 (2021).

44. McLean, C. Y. et al. GREAT improves functional interpretation of cis-regulatory regions. Nature Biotechnology 2010 28:5 28, 495–501 (2010).

45. Baldridge, M. T., King, K. Y., Boles, N. C., Weksberg, D. C. & Goodell, M. A. Quiescent haematopoietic stem cells are activated by IFN-γ in response to chronic infection. Nature 2010 465:7299 465, 793–797 (2010).

46. Kim, T. G. et al. CCCTC-binding factor is essential to the maintenance and quiescence of hematopoietic stem cells in mice. Experimental & Molecular Medicine 2017 49:8 49, e371–e371 (2017).

47. Qi, Q. et al. Dynamic CTCF binding directly mediates interactions among cis-regulatory elements essential for hematopoiesis. Blood 137, 1327–1339 (2021).

48. Pelham-Webb, B. et al. H3K27ac bookmarking promotes rapid post-mitotic activation of the pluripotent stem cell program without impacting 3D chromatin reorganization. Mol Cell 81, 1732–1748.e8 (2021).

49. Demerdash, Y., Kain, B., Essers, M. A. G. & King, K. Y. Yin and Yang: The dual effects of interferons on hematopoiesis. Exp Hematol 96, 1–12 (2021).

50. Essers, M. A. G. et al. IFNalpha activates dormant haematopoietic stem cells in vivo. Nature 458, 904–8 (2009).

51. Baldridge, M. T., King, K. Y., Boles, N. C., Weksberg, D. C. & Goodell, M. A. Quiescent haematopoietic stem cells are activated by IFN-gamma in response to chronic infection. Nature 465, 793–7 (2010).

52. Klimpel, G. R., Fleischmann, W. R. & Klimpel, K. D. Gamma interferon (IFN gamma) and IFN alpha/beta suppress murine myeloid colony formation (CFU-C)N: magnitude of suppression is dependent upon level of colony-stimulating factor (CSF). J Immunol 129, 76–80 (1982).

53. Haas, S. et al. Inflammation-Induced Emergency Megakaryopoiesis Driven by Hematopoietic Stem Cell-like Megakaryocyte Progenitors. Cell Stem Cell 17, 422–434 (2015).

54. Matatall, K. A., Shen, C.-C., Challen, G. A. & King, K. Y. Type II interferon promotes differentiation of myeloid-biased hematopoietic stem cells. Stem Cells 32, 3023–30 (2014).

