## Supplemental Figures for "The Sin3B chromatin modifier restricts cell cycle progression to dictate hematopoietic stem cell differentiation"

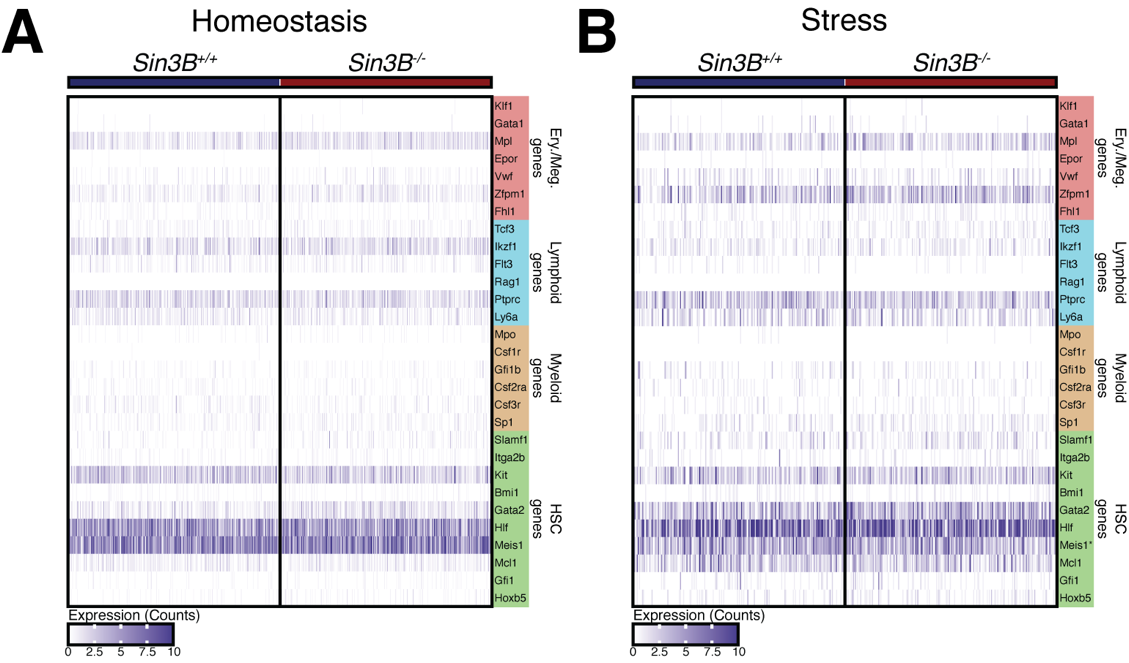

**Supplemental Figure 1. LT-HSCs express lineage-related transcription factors independent of Sin3B status.
A.** Transcription factor expression for specific lineages in LT-HSCs from wild-type or Sin3BH-/- mice at homeostasis. No significant differences were found. Expression from normalized counts shown. **B.** Expression of same transcription factors in A from Sin3B+/+ and Sin3B-/- LT-HSCs. Only Meis1, an HSC stemness gene, was shown to be slightly upregulated in Sin3B-/- LT-HSCs. Other differentiation-related transcription factors were normally expressed. Expression from normalized counts shown.

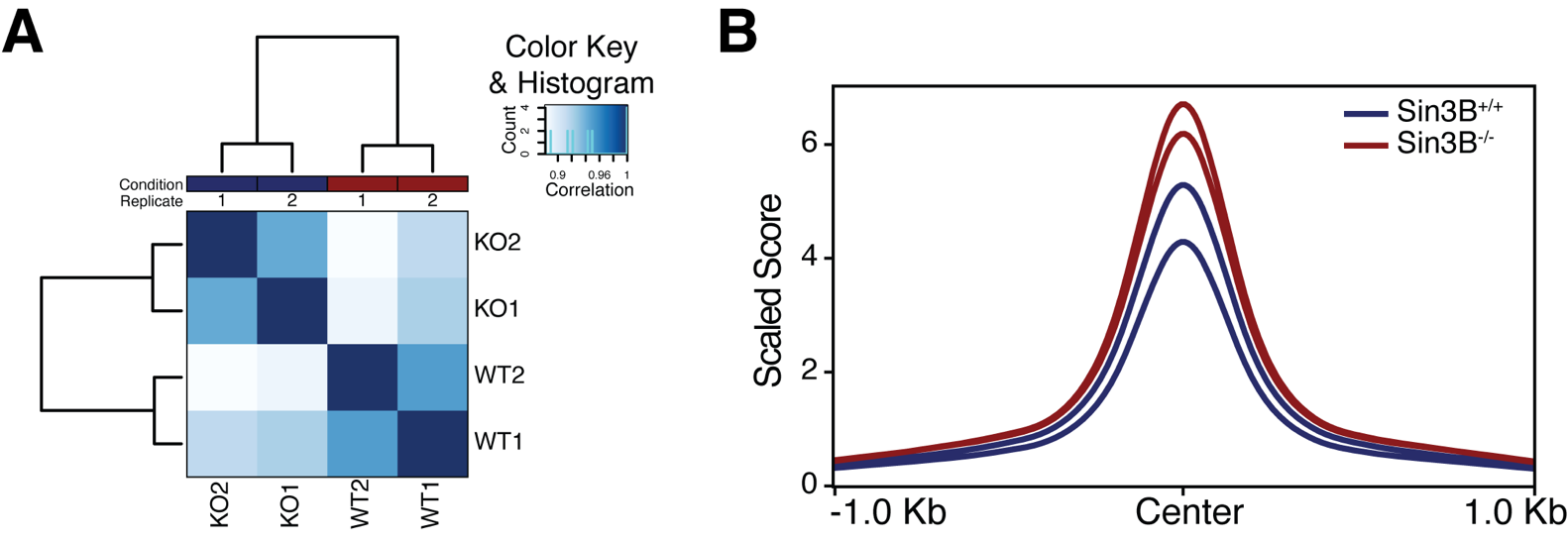
**Supplemental Figure 2. Loss of Sin3B in LT-HSCs results in aberrant chromatin accessibility.**
ATAC-Seq was performed on LT-HSCs isolated via FACS analysis from Sin3B+/+ and Sin3BH-/- mice. **A.** Correlation analysis reveals that wild-type samples (WT1, WT2) cluster closely to one another, and are distinct from Sin3B-/- samples (KO1, KO2). **B.** Analysis of all peaks from samples shows an increase in the level of accessibility in Sin3B-/- LT-HSCs. Each duplicate sample was pooled from 2 mice of the indicated genotype for a total of 8 mice.

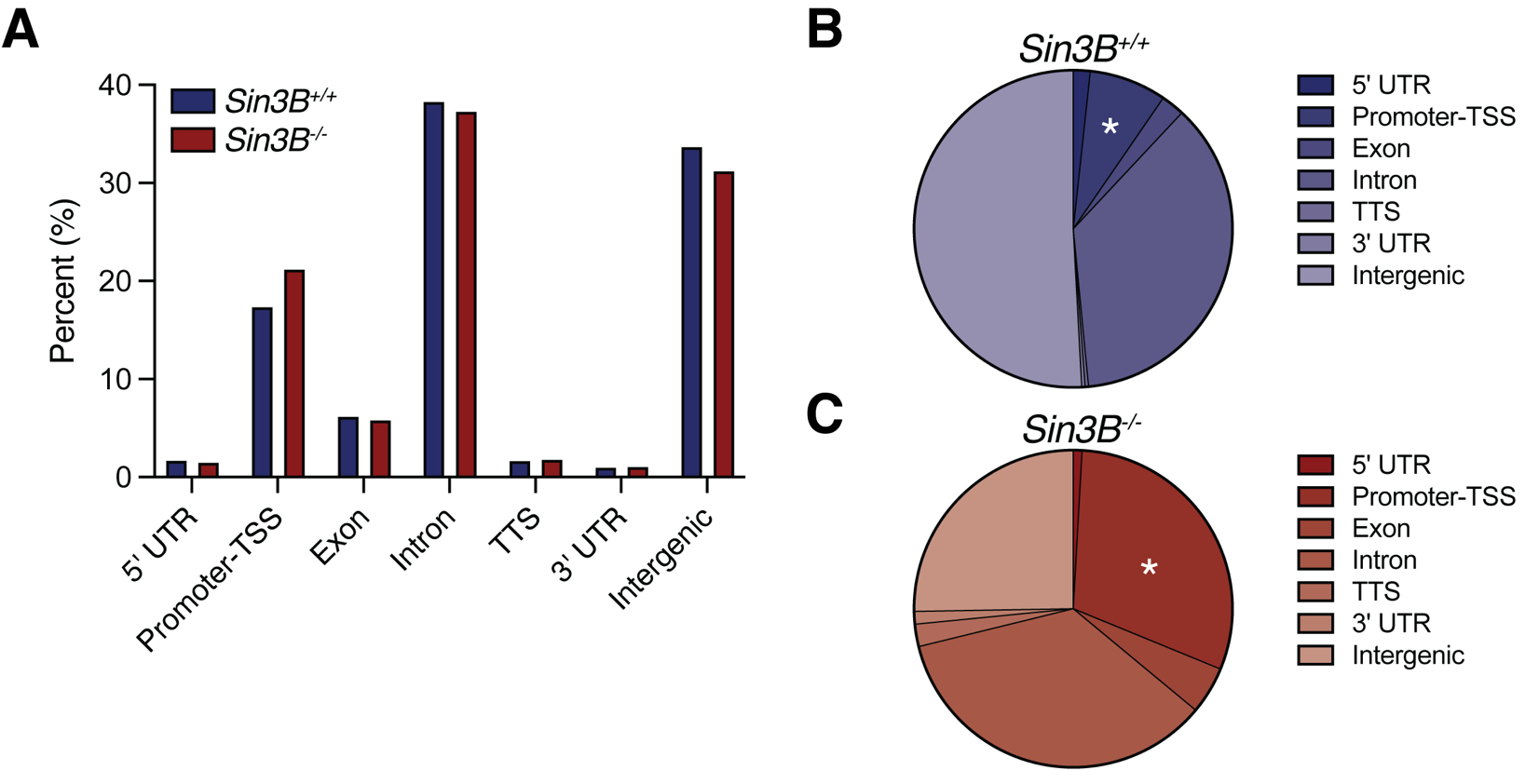

**Supplemental Figure 3. Sin3B is required for promoter silencing.**

**A.** HOMER annotations of all ATAC-Seq data recovered by genotype. **B.** Annotations and proportions of peaks differentially accessible in Sin3B^F/F^ LT-HSCs. **C**. Annotations and proportions of peaks differentially accessible in Sin3B^-/-^ LT-HSCs.
